# Comparison candidate Tick-borne encephalitis virus vaccines based on mRNA, adenovirus serotype 25 and chimera of YFV vaccine strain

**DOI:** 10.1101/2025.01.16.633390

**Authors:** Nadezhda A. Kuznetsova, Vladimir A. Gushchin, Elizaveta V. Marchuk, Andrei E. Siniavin, Denis A. Kleymenov, Maria Nikiforova, Olga V. Zubkova, Olga Popova, Irina V. Vavilova, Tatiana A. Ozharovskaia, Elena P. Mazunina, Evgeniia N. Bykonia, Egor Y. Bogdanov, Anastasia N. Zolotar, Elena V. Shidlovskaya, Evgeny V. Usachev, Olga V. Usacheva, Alexey G. Shatalov, Vladimir I. Zlobin, Denis Y. Logunov, Alexander L. Gintsburg

## Abstract

**Background:** Despite the availability of several licensed inactivated vaccines, the development of new vaccines against the Tick-borne encephalitis virus (TBEV) remains an important task, especially in countries endemic to this pathogen. The risk of infection with TBEV increases every year because of increase in the number of ticks, the emergence of tick carriers into new territories, and active human activity in areas of TBEV natural foci. Annual reports of vaccination failures have prompted us to search for approaches to creating a new vaccine.

**Objectives:** In our study, we used the same PrM and E antigens to produce three vaccine candidates against tick-borne encephalitis (TBE): 1) a live attenuated YFV 17DD-UN vaccine strain, 2) recombinant adenovirus (rAd) vectors, and 3) mRNA encapsulated in lipid nanoparticles (LNP).

**Methods:** We generated and assessed the immunogenicity and protective efficacy of three candidate TBEV vaccines based on mRNA, simian adenovirus type 25, and a chimera of the YFV 17DD-UN attenuated strain.

**Results:** Analysis of the virus-neutralizing titers in the blood sera of immunized mice revealed a statistically significant difference among the three candidate vaccines. The immunogenicity and protective efficacy of the candidate mRNA-LNP vaccine were found to be higher than those of the other two vaccines.

**Conclusions:** Based on the results of our study, it can be concluded that the mRNA-based platform is more promising for the creation of a vaccine against TBEV.

## 1. Introduction

According to WHO data, several thousand cases of tick-borne encephalitis (TBE) are reported each year worldwide [1]. The disease is caused by tick-borne encephalitis virus (TBEV), which belongs to the Flaviviridae family and the Orthoflavivirus genus. In addition to TBEV, the genus Flavivirus includes other arboviruses pathogenic to human such as yellow fever virus (YFV), Japanese encephalitis virus (JEV), dengue virus (DENV), Zika virus (ZIKV) and West Nile virus (WNV) [2]. According to the results of phylogenetic analysis the five subtypes of TBEV were described: European (Eu), Siberian (Sib) and Far-Eastern (FE), Baikalian (Bkl) and new subtype Himalayan (Him) [3,4]. TBEV affects the central nervous system (CNS), leading to neurological consequences of varying severity depending on the TBEV subtype [2,5,6].

In addition to protection against tick bites, vaccination is a good way to prevent disease development, as there is no effective treatment yet [7]. All currently existing licensed vaccines are inactivated and based on the European (strains Neudoerfl and K23 [8,9]) or Far-Eastern (strains Sofjin, 205 [10] and Senzhang [11]) subtypes of TBEV. Although they are quite effective and safe, they have complex immunization schedules which include several doses for primary immunization with necessity for subsequent regular booster vaccinations to maintain intense specific immunity. Meanwhile, it is known that a single vaccination with a live vaccine against yellow fever (also a member of the flavivirus family) provides virtually lifelong immunity [12]. This allows to expect that a vaccine with greater immunogenicity and duration of protection may be created in the future against tick-borne encephalitis.

Study of population immunity in different regions of Russia, including those with a high epidemiological risk of TBE, demonstrated its insufficient level. Most published studies reported vaccination coverage of less than 50% in endemic regions such as the Republic of Altai, Chelyabinsk Region, Zabaykalsky Region, Kemerovo Region, and Kurgan Region [13–15]. The results of the study of population immunity revealed insufficient vaccination coverage of all age groups of study participants, as well as a low proportion of people with protective titers of antibodies to the TBEV among the vaccinated population in all regions of Russia. Thus, the development of new vaccines may provide an opportunity to overcome the shortcomings of existing inactivated TBEV vaccines [16].

Approaches to the development and production of vaccines have evolved over the years, and today several technologies and platforms for their creation already exist, including inactivated, toxoid, live attenuated vaccines, vaccines based on virus-like particles, synthetic peptides, polysaccharides and polysaccharide conjugates, viral vectors vaccines, nucleic acid (DNA and mRNA) vaccines and bacterial vector/synthetic antigen presenting cells [17]. The experience of developing vaccines during the COVID-19 pandemic has shown that platform-based technologies are the best way to respond to emerging infectious diseases. Using a platform approach allows to receive new vaccines quickly, thereby ensuring the safe delivery of the gene encoding the target antigen into the human body [18].

The main components of vaccines are antigens that are obtained directly from the pathogen or biotechnologically [17]. The structure of all flaviviruses, including TBEV, is similar and consists of envelope (formed by membrane (M) and envelope (E) proteins) and nucleocapsid (formed by capsid (C) protein and the RNA genome) [19–22]. In addition to structural proteins, the viral RNA encodes seven non-structural proteins (NS1, NS2A, NS2B, NS3, NS4A, NS4B, NS5) [21], [22]. Antibodies against all large and small TBEV nonstructural proteins can be found in sera and cerebrospinal fluids of patients with TBE [23]. Thus, non-structural proteins stimulate the development of an immune response and may be used as antigens for the creation of vaccines. The E protein of TBEV is a surface glycoprotein, which interacts with attachment factors and/or receptors at the plasma membrane of target cells and promotes fusion of the viral and endosomal membrane, thereby inducing neutralizing antibodies production [19,21]. The precursor-M (prM) protein regulates rearrangement of E proteins on the viral surface leading to transition from immature to mature virions [19,24]. Thus, PrM and E proteins are considered the main antigens and preferred for vaccine development not only against TBEV, but also against other flaviviruses.

In our study, we chose three approaches to PrM/E antigens delivery and candidate vaccine creation: based on a live attenuated YFV 17DD-UN strain (as a genetic backbone for developing vaccines against other viruses), recombinant adenovirus (rAd) vectors (a popular tool for gene delivery into mammalian cells, particularly in the development of vaccine candidates) and mRNA encapsulated in lipid nanoparticles (LNP) (a promising platform of mRNA vaccine development).

Despite the fact that the vaccine against the yellow fever virus was developed 85 years ago, it remains highly effective: studies show that immunity lasts for at least 35 years after just one dose, and antibodies can be detected up to 40 years, which explains the interest in using the genome of live attenuated 17D vaccine strain as the genetic basis for vaccines against other flaviviruses, including tick-borne encephalitis virus. Since 2012, two yellow fever virus-based vaccines have been approved to prevent Japanese encephalitis and dengue [25,26]. Chimeric replication-competent tick-borne encephalitis virus vaccine candidates, including Langat/DENV-4, TBEV/DENV-4, and TBEV/JEV (ChinTBEV), were previously shown to have lower neurovirulence in mice and non-human primates compared with their parent viruses. [27–29]. In our previous work [30], we generated and characterized a chimera of TBEV European subtype (EU) and YFV 17DD-UN strain in which PrM/E genes of YFV 17DD-UN were replaced by PrM/E of TBEV European subtype (EU). During immunization of mice with TBEV (EU)/YFV 17DD-UN the formation of neutralizing antibodies was observed early as 2 weeks after a single dose of the chimeric virus. Thus, firs candidate to deliver PrM and E antigens was replication-competent attenuated chimeric virus based on YFV (YFV 17DD-UN/TBEV(FE)).

Recombinant adenoviral vectors have a good safety profile and induce high levels of humoral and cellular immune responses. Therefore, they are widely used in the development of vaccines against infectious diseases [31–33]. The experience of vaccination during the COVID-19 pandemic has proven the safety and efficacy of adenovirus-based vector vaccines. Moreover, adenovirus-based vaccine candidates have shown to be effective against flaviviruses. In case of the flavivirus E protein, not only neutralizing antibodies, but also T cell-mediated response have been reported [34]. Immunization with adenoviral vectors Ad5-prM-E and Ad4-prM-E have shown a general development of T-cell immune responses [35]. However, there are currently no studies of adenoviral vectors carrying PrM and E genes, therefore, research in this area is relevant. Thus, second candidate to deliver PrM and E antigens was replication-incompetents adenovirus serotype 25 (SAd25-TBEV(FE))

Recently, there has been growing interest in the development of mRNA-based vaccines; such mRNA are not integrated into the genome, they are characterized by prolonged antigen expression, incapable to reversion to pathogenicity, do not cause vector-specific humoral immune reactions, and moreover, mRNA does not need to overcome the nuclear membrane for protein expression that allows transfection of non-dividing or slowly dividing cells such as dendritic cells [36,37]. To protect mRNA from degradation by host nucleases mRNA is encapsulated within a lipid nanoparticle (LNP). During last 10 years several effective mRNA-LNP vaccines were generated against many viruses, including influenza virus [38], HIV [39], rabies virus [40], chikungunya virus [41], human cytomegalovirus [42], flaviviruses DENV [43], ZIKV [44], Powassan [45] and SARS-CoV-2 [46]. In addition, we earlier have shown the effectivity of multivalent mRNA vaccines against seasonal flu [47] and combined mRNA vaccine against influenza and SARS-CoV-2 [48]. As for flaviviruses, it has been demonstrated that cells transfection by mRNA-LNP coding structural proteins prM/E results in the secretion of subviral particles (SVPs) which have similar antigenic features with virus virions and stimulate neutralizing antibodies formation [49]. Therefore, mRNA-LNP platform is a promising approach to creating a vaccine against TBEV, with the ability to quickly updates and combine the antigenic composition if necessary. Thus, third candidate to deliver PrM and E antigens was mRNA-LNP platform (mRNA-TBEV(FE)-LNP).

During our study we generated three candidate TBEV vaccines based on the same Far-Eastern sub-type strain Sofjin. We used three approaches for delivering the PrM and E genes (the major TBEV antigen): mRNA-LNP, recombinant simian adenovirus type 25 and chimera of the YFV 17DD-UN with TBEV genes inserts. Then we evaluated the immunogenicity and protective efficacy of the generated candidate TBEV vaccine-candidates.

## 2. Materials and Methods

### 2.1. Scheme generation of a three candidate vaccines

Developing three candidate vaccines we used Far-Eastern subtype TBEV strain Sofjin (Access ID GenBank KC806252.1). In this study, we used the Sofjin strain stored in the State Virus Collection (SVC) No. 972, the strain was originally isolated by L. A. Zilber from the brain of a person with encephalitis in 1937 (deposited in the collection as № 1). At the first stage, a lyophilized aliquot of the virus SVC No. 972 was used for one passage in the brains of newborn mice. The brain suspension was then used for genome-wide sequencing and Spev cells culture. According to the results of high-throughput sequencing, the sequence was identical to the GenBank ID KC806252.1. TBEV was handled and cultured under biosafety level 3 (BSL-3) conditions.

Virus RNA was used as template for receiving cDNA with High-Capacity RNA-to-cDNA™ Kit (Thermo Fisher Scientific, USA). Oligonucleotide primer for PrM/E cDNA synthesis was as follows: cDNE-TBEV 5’-GTG TCC ACA GCA CAG CCA ACA TC-3’. Then, cDNA was used for amplification of PCR-fragment (size 1,9 kb) with a 2× Platinum SuperFi Green MasterMix kit (Thermo Fisher Scientific, USA). For PCR, primers with the following structure were selected: PrME-For 5’-TGC TGG TTG TTG TCC TGT TGG GA*-*3’ and PrME-Rev 5’-ACC GCC AAG AAC TGT GTG CAG CG-3’ (Figure S1).

Amplification was carried out in accordance with the manufacturer’s instructions. Oligonucleotide primers for synthesis cDNA, amplification of PCR-fragment and constructing plasmids were selected using the SnapGene program (Version 7.2). The identity of the coding sequences was confirmed by Sanger sequencing. Depending on viral vector after one or several passages the presence of the recombinanat viruses was confirmed using high-throughput sequencing.

In the Figure 1 is presented the scheme generation of a chimeric replication-competent attenuated virus YFV 17DD-UN/TBEV(FE), a recombinant replication-defective adenovirus SAd25-TBEV(FE) and mRNA-TBEV(FE)-LNP.

**Figure 1.**
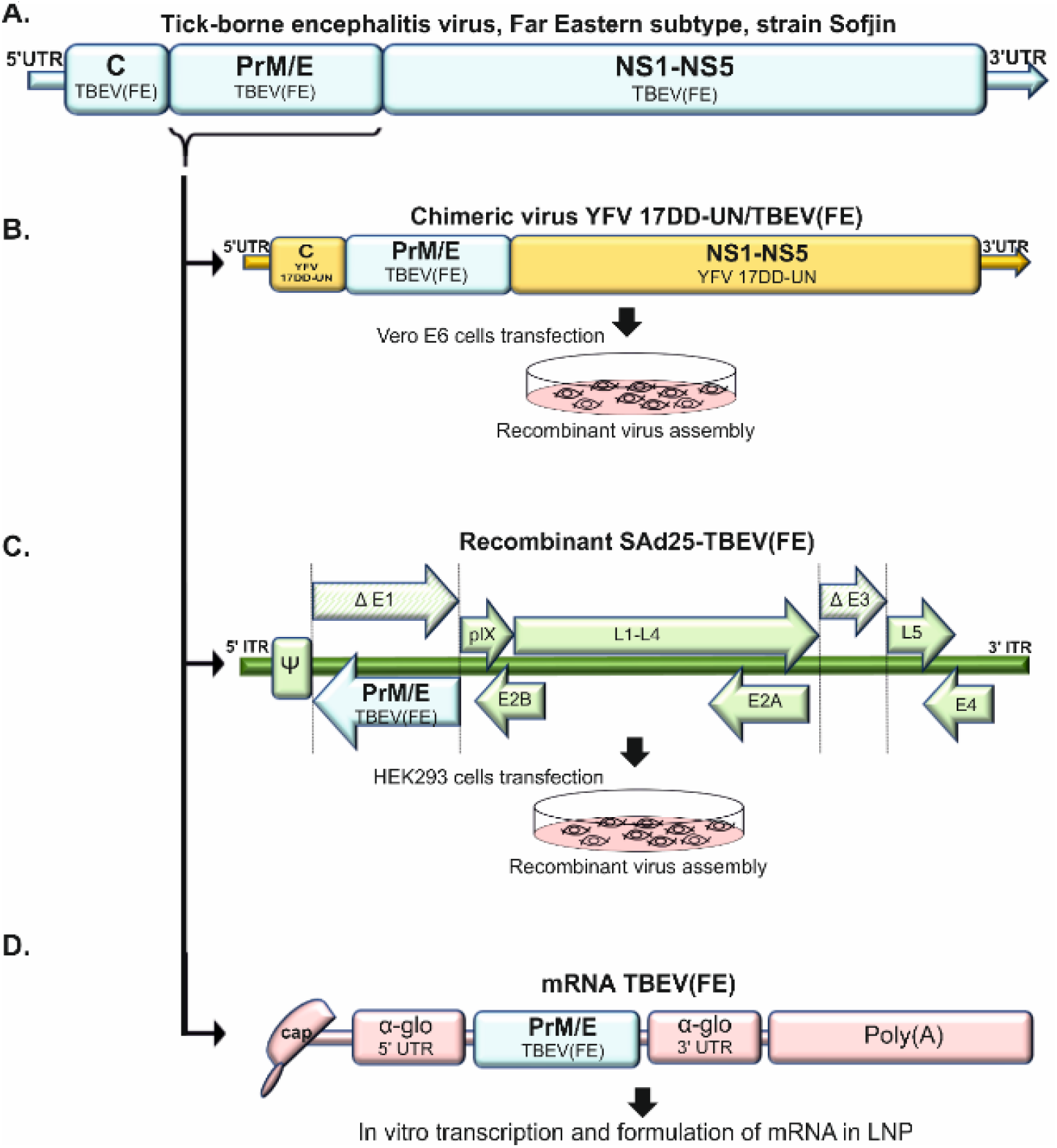
The schematic representation of the design of chimeric virus YFV 17DD-UN/TBEV(FE), a recombinant SAd25-TBEV(FE) and mRNA-TBEV(FE)-LNP used in this study. (a) – The genome scheme of TBEV(FE). (b) - The genome scheme of YFV 17DD-UN/TBEV(FE). (c) – The genome scheme of SAd25-TBEV(FE). (d) – The scheme of mRNA-TBEV(FE)-LNP.

### 2.2. Generation of a chimeric replication-competent attenuated virus YFV 17DD-UN/TBEV(FE)

To generate YFV 17DD-UN/TBEV(FE) chimeric virus based on the YFV 17DD-UN (Access ID GenBank OP796361.1) with PrM/E TBEV genes inserts we used the pSMART BAC plasmid from the Copy-Right v2.0 Bac Cloning kit (Lucigen, USA). We carried all molecular cloning out using the *E. coli* strain Top10. Plasmid construction and assembly were conducted with a Gibson Assembly Ultra MasterMix kit (Codex DNA, USA) following the manufacturer’s instructions.

To assemble hybrid construction, transfection Vero E6 cells was carried out. For this purpose, cells were seeded into 48-well plates (2×10^5^ cells/well) the day before the experiment. Then, cell transfection was performed in OptiMem (Gibco, USA) medium using Lipofectamine 2000 (LF2000; ThermoFisher, USA). Virus replication was confirmed by one-step RT-qPCR and observation the development of virus-induced cytopathic effect (CPE).

Amplification of YFV 17DD-UN/TBEV(FE) chimera was performed using a one-step RT-qPCR technique, as described previously [30]. Oligonucleotide primers and probes for the non-structural protein NS5 gene of the YFV 17DD-UN (backbone) and E gene TBEV (insert) were as follows: YFV forward primer NS5F 5’-GCG GTA TCT TGA GTT TGA GG-3’, reverse primer NS5R 5’-AGG TCT CTG ATC ACA TAT CCT AG-3’, probe NS5TM 5’-FAM-AGC CAA TGC CTTC CAC TCC TCC TC-BHQ1-3’ [30] and TBEV(FE) forward primer ChTBE-Fe-for 5’-GTC AAA GTA GAG CCG CAT AC-3’, reverse primer ChTBE-Fe-rev 5’-T TCT CCG AGG AAG CCG TGA A-3’, probe ChTBE-Fe-Z 5’-R6G-CGT CGC TGC TAA TGA GAC TCA CAG TGG-BHQ1-3’ (this study). To confirm production of PreM and E TBEV antigen virus-containing liquid was examined using enzyme immunoassay (ELISA).

To determine the virus titer, Vero E6 cells were seeded into 24-well plates (2.5×10^5^cells/well) the day before the experiment. Next, various dilutions of the virus were added to the cell monolayer. After virus adsorption, medium was removed, and cells were overlaid with 0.3% Noble Agar (Sigma, USA) solution in DMEM (1:1). Plates were incubated for 4-5 days at 37^0^C and 5% CO_2_. Thereafter, cells were fixed with 5% PFA solution and stained with 1% crystal violet solution. We characterized the titer of the virus in PFU/ml (plaque-forming units) [50].

### 2.3. Generation of a recombinant replication-defective adenovirus SAd25-TBEV(FE)

The recombinant replication-defective vector based on simian adenovirus type 25 (SAd25) containing the TBEV PrM and E genes was obtained by homologous recombination in E. coli BJ5183 cells. Bacterial cells were transformed by electroporation according to the manufacturer’s instructions using a MicroPulser (Bio-Rad, Hercules, USA). The resulting plasmid (pSAd25-TBEV(FE)) was analyzed by PCR, restriction mapping and high-throughput sequencing following standard protocols [30], [51]. In pSAd25-TBEV(FE) the deleted E1 region of the adenoviral genome was replaced by a cassette containing the target gene (CMV promoter, PrM and E genes of TBEV from the same strain used to generate the chimeric YFV 17DD-UN/TBEV(FE), along with a polyadenylation signal). To increase the packaging capacity, the E3 region of the adenoviral genome was also deleted.

The pSAd25-TBEV(FE) plasmid was used to generate the recombinant adenovirus SAd25-TBEV(FE). HEK293 cells were seeded in 24-well culture plates and incubated overnight to reach 80% confluence. To remove the bacterial component, plasmid DNA was digested with restriction endonucleases PacI and SspI and then transfected into HEK293 cells using Lipofectamine 2000 reagent (Thermo Fisher Scientific, USA) according to the manufacturer’s instructions. After visual detection of viral cytopathic effect (CPE) using a CKX41 inverted microscope (Olympus, Japan), the cells in the culture medium were subjected to three freeze-thaw cycles.

Recombinant simian adenovirus was accumulated in HEK293 cell culture. HEK293 cells were seeded in 10 cm diameter culture dishes at a density of 15–17×106 cells per dish. The next day, a 65–75% confluent cell monolayer was infected with recombinant SAd25-TBEV(FE) at a dose of 10^7^ PFU per dish. Two days later, when 90–100% CPE was achieved, the infected cells were collected, concentrated by low-speed centrifugation, resuspended in buffer (0.01 M Tris-HCl pH 8.0, 0.01 M NaCl, 5 mM EDTA), and subjected to three freeze-thaw cycles to disrupt the cellular and nuclear membranes and release the virus from the cells. Cell lysates were centrifuged at 5000 rpm for 10 minutes at room temperature, and the precipitate was removed. The recombinant adenovirus was then purified by double ultracentrifugation in a CsCl gradient (both step and equilibrium gradients) using an Optima XPN-90 ultracentrifuge (Beckman Coulter Inc., USA).

The purity and identity of SAd25-TBEV(FE) were confirmed by PCR and whole-genome sequencing. The titer of the purified virus was determined using TCID50 assay on a HEK293 cells [52]. The results were recorded on the 10th–12th day after cell transduction. The number of virus particles was determined using reagents from the Pico488 dsDNA quantification kit (Lumiprobe, USA) in accordance with the manufacturer’s instructions.

SDS-PAGE was performed in 12% polyacrylamide gel according to the Laemmli method [53]. Precision Plus Protein™ was used as a molecular weight standard. Particle size was determined using a Zetasizer Nano ZS nanosizer (dynamic light scattering, DLS). Adenovirus preparations were diluted to a concentration of 105 v.p./ml in filtered (PES filter, pore size 0.22 μm) milliQ water. Then 1 ml of particles was transferred to a 10×10×45 mm cuvette for measurement and triplicate measurements were performed. Particle size was calculated using Zetasizer Software.

For expression analysis, 3 cm diameter culture dishes with 80-90% monolayer of VeroB cells were inoculated with SAd25-TBEV(FE) and the control SAd25-EGFP virus (expressing reporter protein EGFP gene). After 72 h, the cells and medium were collected and analyzed by ELISA.

### 2.4. Generation of a mRNA-TBEV(FE)-LNP

The development of mRNA platform for efficient and long-term expression of the gene of interest, including DNA cloning procedures, the production and purification of plasmid DNA, in vitro transcription (IVT) and formulation of mRNA in LNP were described previously [54]. The pJAZZ-OK-based linear bacterial plasmid (Lucigen, USA) with coding regions of PrM/E was used as templates for mRNAs production. The pDNA for IVT were isolated from the *E*.*coli* BigEasy™-TSA™ Electrocompetent Cells (Lucigen, USA) using the Plasmid Maxi Kit (QIAGEN). The pDNA was digested using BsmBI-v2 restriction endonuclease (NEB), followed by purification of the product by phenol-chloroform extraction and ethanol precipitation. IVT was carried out as described earlier [50]. Prepared a 100-μl reaction mixture, contained 3 μg of DNA template, 3 μl T7 RNA polymerase (Biolabmix) and T7 10X Buffer (TriLink), 4 mM trinucleotide cap 1 analog (3′-OMe-m7G)-5′-ppp-5′-(2′-OMeA)pG (Biolabmix), 5 mM m1ΨTP (Biolabmix) replacing UTP, and 5 mM GTP, ATP and CTP. After 2 h incubation at 37°C, 6 μl DNase I (Thermo Fisher ScientifiC, USA) was added for additional 15 min, followed by mRNA precipitation with 2M LiCl (incubation for 1 h in ice and centrifugation for 30 min at 14,000 g, 4°C) and carefully washed with 80% ethanol. RNA integrity was assessed by electrophoresis in 8% denaturing PAGE.

LNP assembly also was described previously [54]. All lipid components were dissolved in ethanol at molar ratios 46.3:9:42.7:1.6 (ionizable lipid:DSPC:cholesterol:PEG-lipid). Acuitas ionizable lipid (ALC-0315) and PEG-lipid (1,2-Dimyristoyl-sn-glycero-3-methoxypolyethylene glycol 2000) were purchased in Cayman Chemical Company. 1,2-distearoyl-sn-glycero-3-phosphocholine (DSPC) and Cholesterol were purchased in AvantiResearch and Merck respectively. The lipid mixture was combined with an acidification buffer of 10 mM sodium citrate (pH 3.0) containing mRNA (0.2 mg/mL) at a volume ratio of 3:1 (aqueous: ethanol) using the NanoAssemblr Ignite device (Precision NanoSystems, Canada). The ratio of ionizable nitrogen atoms in the ionizable lipid to the number of phosphate groups in the mRNA (N:P ratio) was set to 6 for each formulation. Formulations were dialyzed against PBS (pH 7.2) in Slide-A-Lyzer dialysis cassettes (Thermo Fisher Scientific, USA) for at least 24 h. Formulations were passed through a 0.22-μm filter and stored at 4°C (PBS) until use. The diameter and size distribution, zeta potential of the mRNA-LNP were measured using a Zetasizer Nano ZS instrument (Malvern Panalytical, United Kingdom) according to user manual.

The mRNA encapsulation efficiency and concentration were determined by SYBR Green dye (SYBR Green I, Lumiprobe) and were described earlier [47, 54]. The HEK293 cells transfection with PreM/E encoding mRNA was performed as described previously [48, 54]. To confirm production of PreM and E TBEV antigen cell liquid was examined by ELISA.

### 2.5. Dynamic light scattering

The particle size was determined using the Zetasizer Nano ZS nanosizer (Malvern, UK) - using the dynamic light scattering (DLS) method. The DLS method determines the diffusion coefficient of dispersed particles in a liquid after analyzing the correlation function of fluctuations in the intensity of scattered light. Then, the particle radius is calculated from the determined diffusion coefficient.

For the Zetasizer Nano ZS, SAd25-TBEV(FE) adenovirus sample was diluted to a concentration of 7.3 ×10^9^/ml in filtered (PES filter, pore size 0.22 μm) milliQ water. Then 1 ml of the diluted sample was transferred to a measurement cuvette (10×10×45 mm, polystyrene/polystyrene material, Sarstedt, Germany). Then, the particle size was measured three times.

The mRNA-LNP suspension was diluted in water 20 times and loaded into a cuvette for measurement at 25 °C. LNPs in PBS were read using the solvent parameter ‘water’. One measurement consists of three readings and each reading is derived from 13 acquisitions. The DLS data presented for mRNA-LNP preparation are the average value of these three readings.

The results were calculated using Zetasizer Software (Malvern, UK).

### 2.6. The mRNA encapsulation efficiency

The mRNA encapsulation efficiency and concentration were determined by SYBR Green dye (SYBR Green for PCR, Lumiprobe, Moscow, Russia) followed by fluorescence measurement. Briefly, mRNA-LNP samples were diluted with TE buffer (pH 8.0) in the absence or presence of 2% Triton-X-100 in a black 96-well plate. Standard mRNA was serially diluted with TE buffer in the absence or presence of 2% Triton-X-100 to generate standard curves. Then, the plate was incubated for 10 min followed by the addition of SYBR Green dye (100 times diluted) to each well to bind RNA. Fluorescence was measured at 497 nm excitation and 522 nm emission using Varioscan LUX (Thermo Fisher Scientific Inc., Waltham, MA, USA). The concentrations of mRNA after LNP disruption by Triton-X-100 (C total mRNA) and before LNP disruption (C outside mRNA) were determined using corresponding standard curves. The concentration of mRNA loaded into the LNP was determined as the difference between the two concentrations multiplied by the dilution factor of the original sample. Encapsulation efficiency was calculated by the formula: (E%) = (C total mRNA)–(C outside mRNA)]/ (C total mRNA) × 100%.

### 2.7. Enzyme immunoassay (ELISA) to detect PrM/E antigens

Medium or virus-containing liquid were collected and analyzed using the VectoVKE-antigen tick-borne encephalitis virus antigen detection reagent kit (Vector-Best, Russia).

We evaluated ELISA resalts in the μ-Quant device (Bio-Tek Instruments, USA) at a wavelength of 450 nm with the reference wavelength of 620 nm. Calculation of ELISA cut-off value was carried out in accordance with the manufacturer’s instruction by using Formular (1):

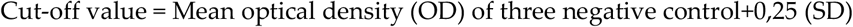

### 2.8. Animal studies

All animal procedures were conducted in accordance with the relevant guidelines for the care and use of laboratory animals and approved by the Local Ethics Commission of the National Research Center for Epidemiology and Microbiology named after Honorary Academician N.F. Gamaleya (protocol # 63, 05 October 2023). In according to the institutional and national guidelines, all experiments were performed in the BSL-2 and−3 facilities.

Mice were purchased from Stolbovaya nursery for laboratory animals (Russia). All animals were housed in separate cages (10 mice in cage) with controlled temperature (20-24°C) and humidity (45-65%). Mice were fed with a balanced rodent diet and water ad libitum during the entire study.

### 2.9. Immunizations

To access the immunogenicity and protectiveness of generated recombinant viruses and mRNA-LNP, BALB/c female mice (6–8 weeks of age) were randomly assigned into groups (n = 10 per group). Depending on the platform used for delivery PrM/E of TBEV (as viruses or mRNA-LNP) mice were injected subcutaneously or intramuscularly 1 or 2 times. Control group included mice injected with phos-phate-buffered saline (PBS) (placebo group, n = 10). Mice health and behavior were monitored once a day till the end of the study. In case of the observation of neurological signs such as paresis mice were sacrificed by cervical dislocation. After 28 days of observation, blood samples were collected from the tail vein. Blood serum was collected to determine the level of neutralizing antibodies.

### 2.10. TBEV challenge

To analyze the protective properties of recombinant viruses and mRNA-LNP, we infected immunized mice with TBEV strain Sofjin. After immunization animals all groups were intraperitoneally inoculated by TBEV with a dose 100 LD_50_. The LD_50_ was calculated using the Reed and Muench method [55]. Mice health and behavior were monitored once a day during one week after TBEV challenge and then twice a day till the end of the study. In case of the observation of neurological signs such as paresis mice were sacrificed by cervical dislocation. After 20 days of observation, surviving animals were sacrificed by cervical dislocation.

### 2.11. Evaluation of serum neutralizing activity after immunization

Different dilutions of serum from immunized animals were mixed in equal proportion with 100TCID_50_ of TBEV strain Sofjin and incubated for 1 hour at 37^0^C. After incubation, serum-virus mix was transferred into a 96-well plate to Spev cells (pig kidney embryonic cells). Plates were incubated at 37^0^C and 5% CO_2_ for 4 days. The last dilution of the serum that showed full protection against the virus-induced cytopathic effect (CPE) was taken as the viral neutralizing titer.

### 2.12. Statistical Analyses

In the mouse experiments, body weight change between groups was performed using two-way ANOVA. All differences were considered statistically significant at *p* ≤ 0.05. To interpret Kaplan-Meier survival analysis was used the log-rank (Mantel-Cox) test. To compare level viral neutralizing antibodies was used the Kruskal-Wallis test followed by Dunn multiple comparison post-hoc test. All Figures were generated by using GraphPad software, 8th edition (GraphPad Software, San Diego, CA, USA).

## 3. Results

### 3.1. Generation of a chimeric replication-competent attenuated virus YFV 17DD-UN/TBEV(FE)

In our previous study [30], we generated chimeric YFV 17DD-UN/TBEV(Eu) virus by using infectious subgenomic amplicon (ISA) approach. Here we made chimeric YFV 17DD-UN/TBEV(FE) by using bacterial artificial chromosomes. For this purpose, an infectious cDNA backbone YFV 17DD-UN was cloned into pSMART BAC plasmid, then cell was transfected by artificial chromosomes with subsequent evaluation virus-induced cytopathic effect (CPE) and confirmation virus replication by RT-qPCR. CPE and virus replication was observed in Vero E6 cells both after transfection and after virus passages. After several passages in Vero E6 cells, the presence of the full YFV 17DD-UN genome has been confirmed by high-throughput sequencing. In accordance with a comparative analysis of the complete genomes of YFV 17DD-UN (Access ID GenBank OP796361.1) (generated by ISA) and YFV 17DD-UN (generated as artificial chromosomes) in the presence study no nucleotide substitutions were detected.

Artificial chromosome containing complete genom of YFV 17DD-UN strain was then used as a backbone in the development of the chimeric virus. The structural genes PrM and E of backbone YFV 17DD-UN were replaced by the same genes of TBEV. Plasmid constraction was assembled by Gibson method (Gibson assembly). Sanger sequencing confirmed identity PrM and E of TBEV(FE) in YFV 17DD-UN backbone. The following transfection and passages in Vero E6 cells showed that the construction of the chimeric virus YFV 17DD-UN/TBEV(FE) was successful, which was confirmed by CPE and RT-qPCR. The third and six passages of the hybrid virus were used for assessment genom stability by high-throughput sequencing. In the result of a high-throughput sequencing, a no nucleotide substitutions were detected.

The virus-containing liquid collected from Vero E6 cells was also examined using enzyme immunoassay (ELISA), which confirmed the production of PreM and E TBEV antigen proteins by a chimeric virus (Table 1).

**Table 1.** Production of PreM and E TBEV antigen evaluated by ELISA.

|  | Optical density (mean) |
| --- | --- |
| Negative control (buffer for dilution) | 0,053 |
| Cut-off value | 0,30 |
| YFV 17DD-UN/TBEV(FE) (culture medium) | 2,609 |
| YFV 17DD-UN/TBEV(FE) (cell lysate) | 3,908 |
| SAd25-TBEV(FE) (culture medium) | 3,598 |
| SAd25-TBEV(FE) (cell lysate) | 4,000 |
| SAd25-EGFP (culture medium)" | 0,051 |
| SAd25-EGFP (cell lysate) | 0,057 |
| mRNA-PrM/E-TBEV(FE)(culture medium) | 3,653 |
| mRNA-PrM/E-TBEV(FE)(cell lysate) | 4,000 |
| HEK293 (culture medium) | 0,049 |
| HEK293 (cell lysate) | 0,060 |
| Vero E6 (culture medium) | 0,055 |
| Vero E6 (cell lysate) | 0,056 |
| Positive control | 3,106 |

### 3.2. Generation of a recombinant replication-defective adenovirus SAd25-TBEV(FE)

Previously, we developed a technological platform based on SAd25 [51]. Here, we used the same approach to obtain a recombinant simian adenovirus containing the TBEV PrM and E genes. One of the important aspects of successfully complementing E1-deleted adenoviruses is the functional interaction of the E1B 55K protein (produced by the trans-complementing cell line) with the E4 34K protein in the viral genome. However, the development of complementing cell lines for different types of replication-defective vectors is a labor-intensive and cumbersome process. Therefore, the availability of non-human adenovirus types capable of replication in cells such as HEK293 is a significant advantage.

To obtain the SAd25-TBEV(FE) virus, HEK293 cells were transfected with the plasmid construct pSAd25-TBEV(FE). We demonstrated that this cell line efficiently assembles recombinant simian adenovirus type 25. The sequence similarity between the Ad5 and SAd25 E1B 55K proteins is approximately 56%. Therefore, it was surprising that we were able to successfully revive replication-defective SAd25 in HEK293 cells.

After virus growth and purification, we measured the particle size and analyzed the protein spectrum. The size of the SAd25-TBEV(FE) virion is 108.23 ± 26.64 nm (Figure 2a). Analysis of the protein spectrum of the recombinant adenovirus SAd25-TBEV(FE) indicates the presence of major capsid proteins—hexon (II), penton base (III), and fiber (IV)—as well as minor proteins (V, VI, VIII, IX) and some core proteins (VII). The molecular weight of SAd25-TBEV(FE) polypeptides was approximately as follows: hexon (pII) – 120-170 kDa; penton base (pIII) – 51-65 kDa; pIIIa and fiber – 60-65 kDa; minor core protein pV ∼ 48 kDa; hexon associated protein pVI – 22-24 kDa (Figure 2b). The expression of the pre-M and E protein genes was determined by ELISA (Table 1).

**Figure 2.**
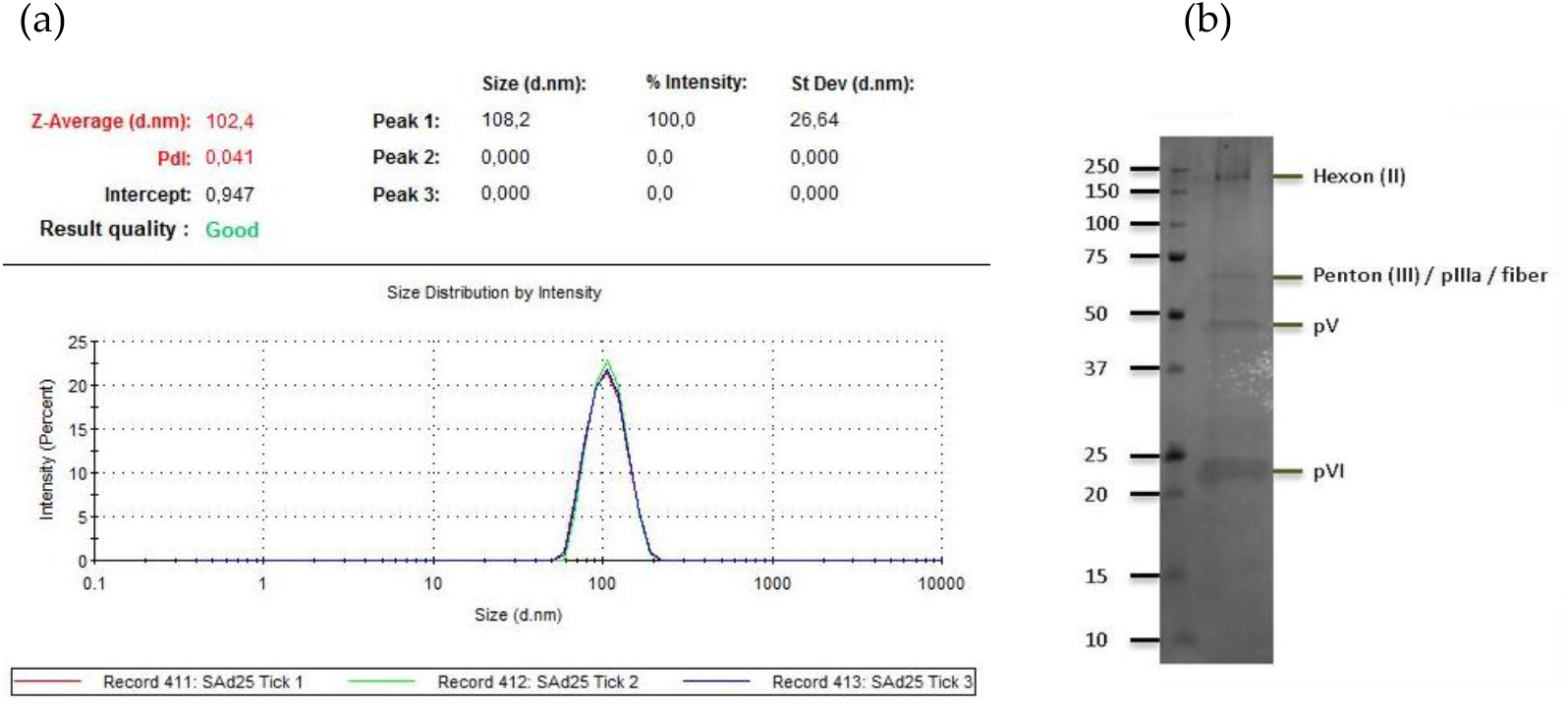
Characteristics of the recombinant simian adenovirus SAd25-TBEV(FE). (a) results of the analysis of the homogeneity of the distribution of recombinant adenoviral particles by size. (b) SDS-PAGE of SAd25-TBEV(FE).

### 3.3 Generation of a mRNA-TBEV(FE)-LNP

To generate mRNA-TBEV(FE)-LNP PrM/E genes of TBEV were synthesized and cloned into the pJAZZ-OK linear bacterial plasmid, as described previously [47, 54]. In vitro synthesized mRNAs included the cap-1 structure at the 5′ end; a 100-nt long poly(A)-tail at the 3′ end; the 5′ and 3′ untranslated regions (UTRs) from the human hemoglobin alpha subunit (HBA1) mRNA.

N1-methylpseudouridines (m1Ψ) were co-transcriptionally incorporated into the mRNA instead of 100% uridines (U). mRNA-LNP formulations were prepared using the microfluidic NanoAssemblr Ignite mixer. The encapsulation efficiency was 90% (SD 1.2%) with a particle size 88,64 nm ± 32,4 nm (Figure 3).

**Figure 3.**
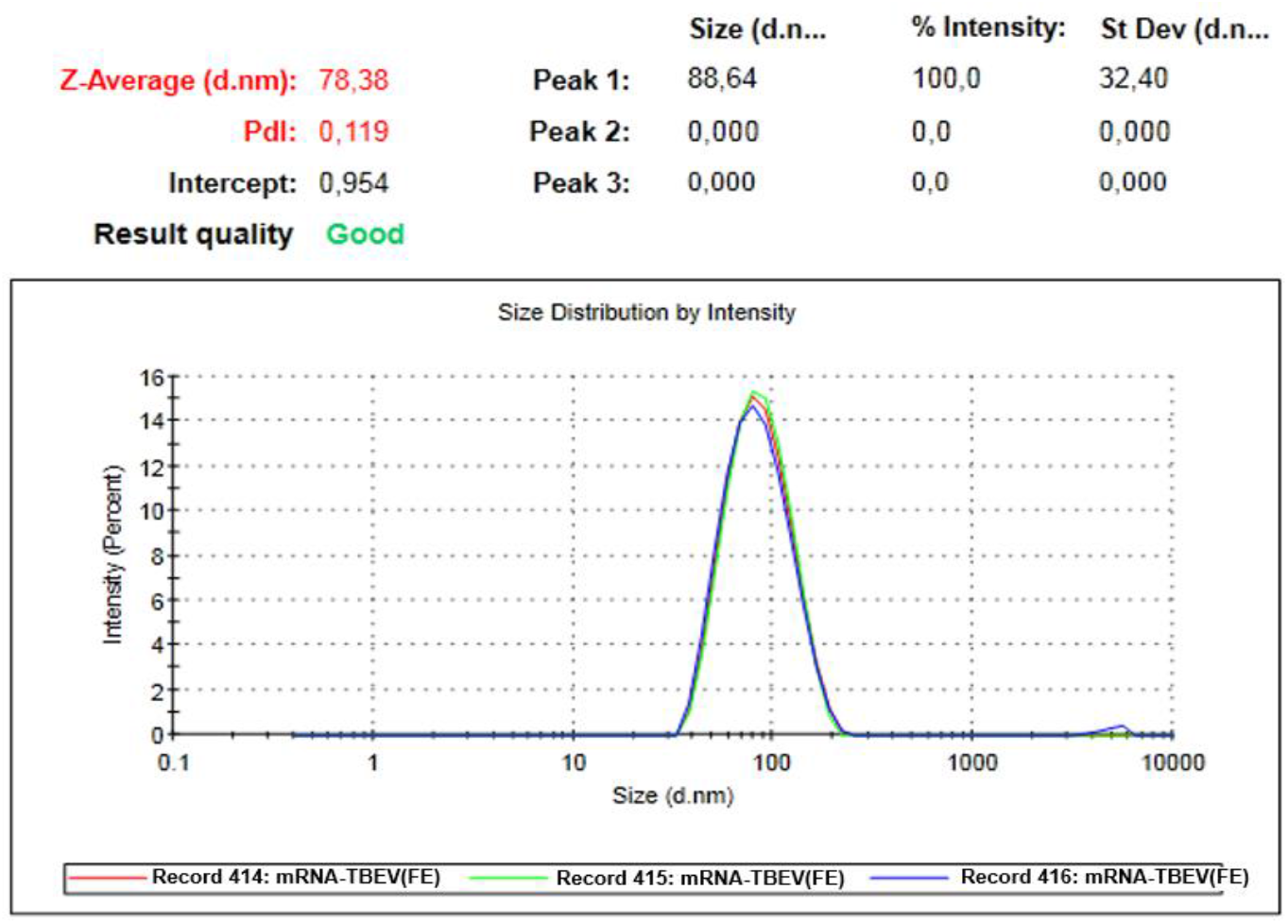
Results of the analysis of the homogeneity of the distribution of mRNA-TBEV(FE)-LNP particles by size.

### 3.4. Estimation of the PrM/E antigens production

An expression of the PrM/E antigens by in vitro synthesized mRNA-TBEV(FE)-LNP, recombinant replication-defective adenovirus SAd25-TBEV(FE) and replication-competent attenuated virus was confirmed by transfection of cultured HEK293 and Vero E6 respectivly. Culture medium and/or cell lysate were then examined in the ELISA (Table 1). To exclude unspecific serological reactions culture medium and cell lysate of HEK293 and Vero E6 were used for ELISA. For the same purpose, culture medium and cell lysate of SAd25-EGFP (SAd25 containing gene of green fluorescent protein (EGFP)) were studied in ELISA.

These results confirmed the translational activity of our synthetic mRNA and the presence of an antigen in generated viruses YFV 17DD-UN/TBEV(FE) and SAd25-TBEV(FE).

### 3.5. Estimation of the immunogenicity of chimeric replication-competent attenuated virus YFV 17DD-UN/TBEV(FE), a recombinant replication-defective adenovirus SAd25-TBEV(FE) and mRNA-TBEV(FE)-LNP

To assess the immunogenicity of generated recombinant YFV 17DD-UN/TBEV(FE) and SAd25-TBEV(FE) viruses and mRNA-TBEV(FE)-LNP, BALB/c mice were subcutaneously inoculated with the YFV 17DD-UN/TBEV(FE) virus at dose of 10^4^ PFU per mouse (YFV 17DD-UN/TBEV(FE) group, n=10), intramuscularly immunized with SAd25-TBEV(FE) at dose of 5×10^10^ viral particles (VP) per mouse (SAd25-TBEV(FE) 2 groups, each group n=10) and intramuscularly immunized with mRNA-TBEV(FE)-LNP at dose of 5 μg per mouse (mRNA-TBEV(FE)-LNP group, n=10). Doses were selected during preliminary experiments. Animals injected with PBS were used as control groups (group Placebo, n=10). Three groups of BALB/c mice received one dose of the YFV 17DD-UN/TBEV(FE), SAd25-TBEV(FE) and placebo, the other two groups received two doses of the SAd25-TBEV(FE) and mRNA-TBEV(FE)-LNP (Figure 4).

**Figure 4.**
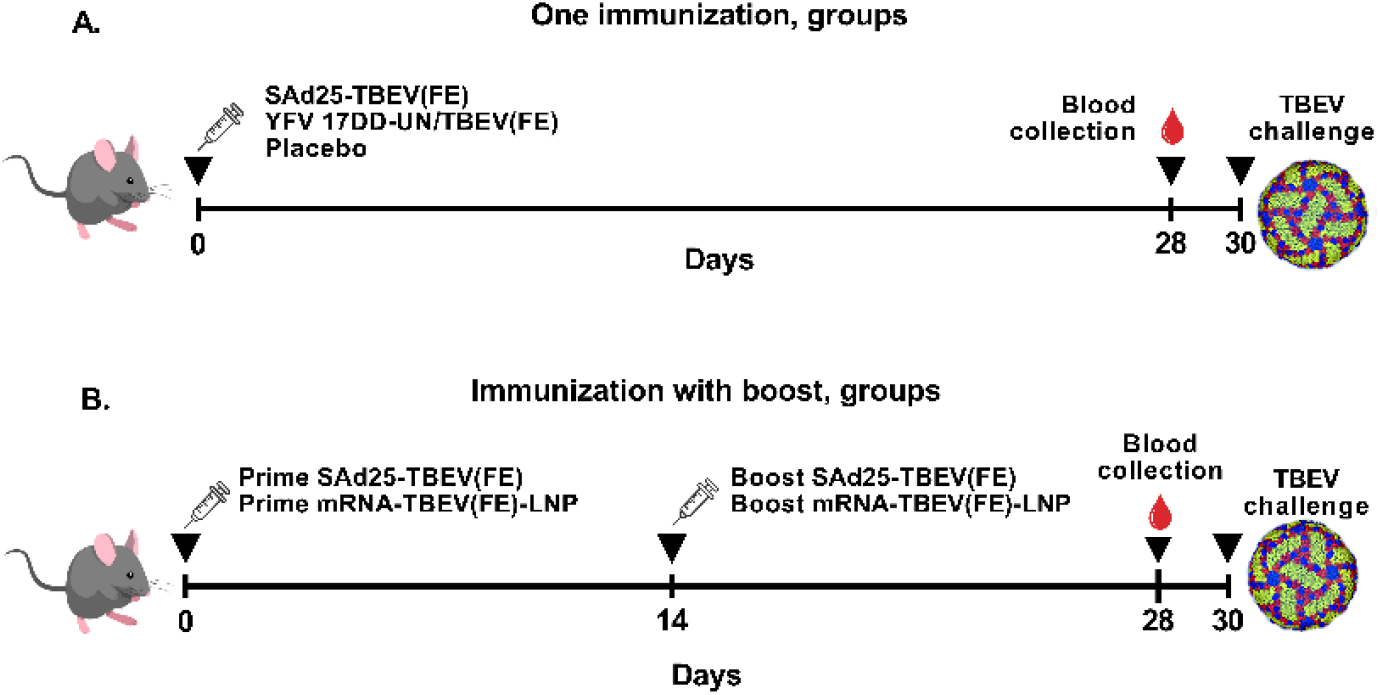
Timeline of immunization and blood sample collection. A – Groups animals with one immunization. B – Groups animals with prime and boost immunization.

After immunization, the animals were observed for 4 weeks. During the experiment, changes in the animals’ body weight and behavioral were assessed. Neurological signs such as paralysis were observed in only 2 animals in the YFV 17DD-UN/TBEV(FE) chimeric virus group. A two-way analysis of variance (ANOVA) revealed no statistically significant differences in changes in body weight (an indicator of animal welfare) either between or within groups.

To assess the neutralizing activity of the serum, blood samples were collected 28 days after immunization. It was found that immunization of mice with 10^4^ PFU of YFV 17DD-UN/TBEV(FE) did not lead to the formation of neutralizing antibodies. Neutralizing antibodies were detected in half of the animals receiving one dose of SAd25 TBEV(FE). Immunization with two doses of SAd25 TBEV(FE) and mRNA-TBEV(FE)-LNP led to the formation of antibodies in all animals in the groups (Figure 5).

**Figure 5.**
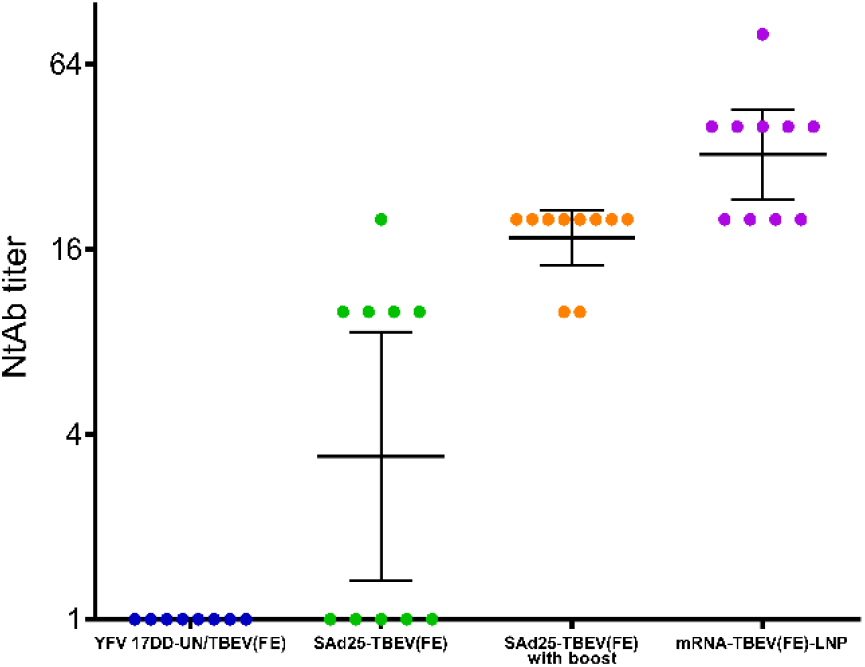
Titer of virus neutralizing antibodies in the blood serum of mice immunized with YFV 17DD-UN/TBEV(FE), SAd25-TBEV(FE) and mRNA-TBEV(FE)-LNP. Each dot represents the serum titer for each animal that protects 100% against CPE.

Analysis of the titer of virus neutralizing antibodies in the blood sera showed a statistically significant difference between all groups. In the mRNA group, the antibody titer was higher than in other groups. The Kruskal-Wallis test was used for calculations, p value is presented in the Table 2.

**Table 2.** Comparison of virus neutralizing antibody titers in the blood sera of mice immunized with YFV 17DD-UN/TBEV(FE), SAd25-TBEV(FE) and mRNA-TBEV(FE)-LNP. P value was calculated using Kruskal-Wallis test.

|  | YFV 17DD-UN/TBEV(FE) | SAd25 TBEV(FE) | SAd25 TBEV(FE) with boost dose | mRNA-PrM/E-TBEV(FE) |
| --- | --- | --- | --- | --- |
| YFV 17DD-UN/TBEV(FE) | - | - | - | - |
| SAd25 TBEV(FE) | >0,9999 | - | - | - |
| SAd25 TBEV(FE) with boost dose | 0,0042** | 0,0987 | - | - |
| mRNA-PrM/E-TBEV(FE) | <0,0001**** | 0,0004*** | 0,6586 | - |

### 3.6. Estimation of the protectivity of the chimeric replication-competent attenuated virus YFV 17DD-UN/TBEV(FE), the recombinant replication-defective adenovirus SAd25-TBEV(FE) and the mRNA-TBEV(FE)-LNP

To assess the ability of the chimeric replication-competent attenuated virus YFV 17DD-UN/TBEV(FE), the recombinant replication-defective adenovirus and SAd25-TBEV(FE) and the mRNA-TBEV(FE)-LNP the to protect animals from lethal TBEV infection, we infected immunized animals with TBEV (strain Sofjin). Experimental challenge of animals with a dose of 100 LD_50_ resulted in 100% survival in the groups mRNA-TBEV(FE)-LNP and in the both SAd25-TBEV(FE) groups. Challenge with TBEV Sofjin at 100 LD_50_ resultedin 85,7% and 0% survival rate in the groups YFV 17DD-UN/TBEV(FE) and Placebo, respectively. Significant differences among the survival curves (p < 0.0001) were showed by logrank (Mantel-Cox) test (Figure 6).

**Figure 6.**
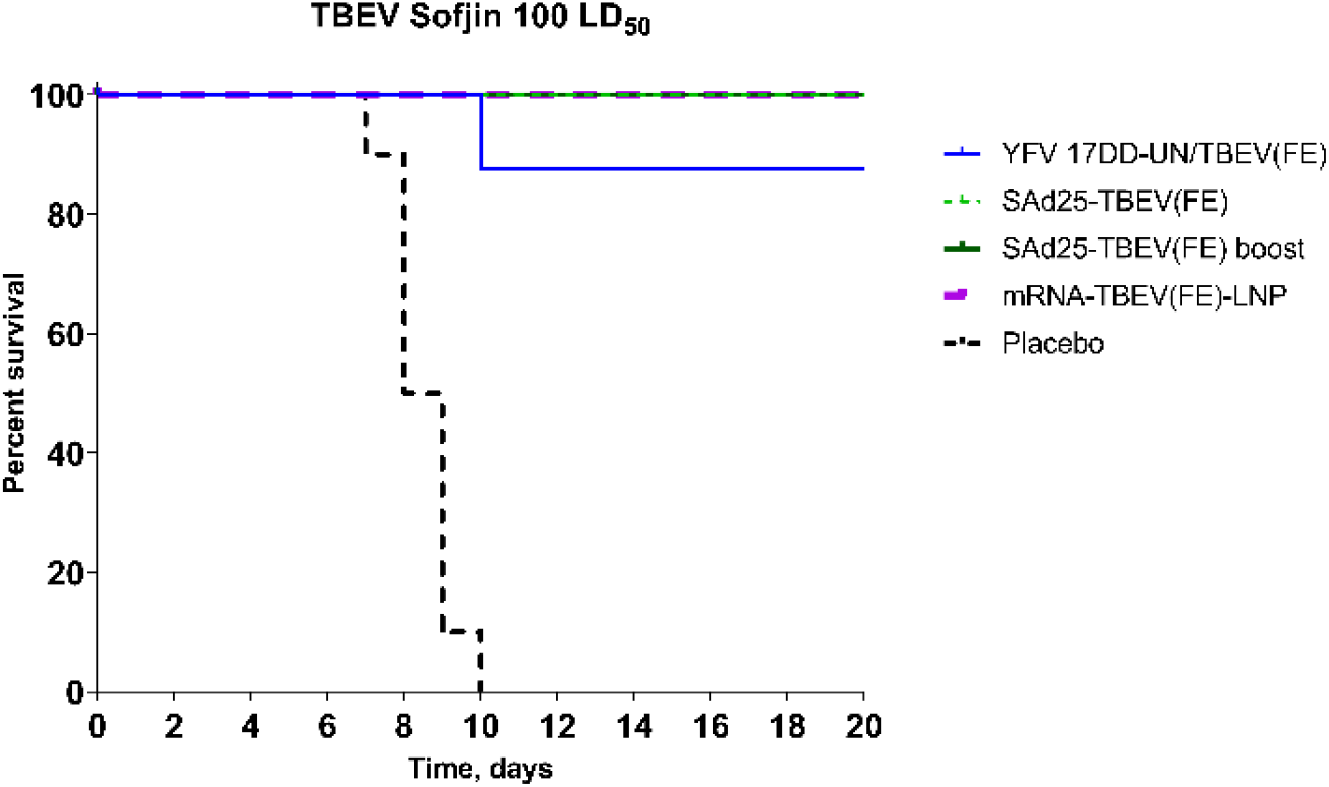
Assessment protectivity of the YFV 17DD-UN/TBEV(FE), the SAd25-TBEV(FE) and the mRNA-TBEV(FE)-LNP. The mortality of animals was observed for 20 days. Kaplan-Meier survival analysis immunized mice, TBEV Sofjin strain challenge by 100 LD_50_.

## 4. Discussion

All currently existing conventional inactivated TBEV vaccines do not provide 100% protection against infection and disease development. TBE cases are reported annually among vaccinated individuals [56–58]. The reasons for the TBEV vaccines failures are currently being studied by different groups of researchers, thereby providing thorough understanding of the immune responses of protection against TBEV [59,60]. Development of improved vaccines and new vaccination strategies may serve as one of the ways to overcome impaired vaccine responsiveness [16].

Conventional vaccine production technologies have guided vaccine development over the past century, effectively protecting against diseases with high rates of disability and mortality. The infrastructure and resources required for these technologies already exist. However, traditional technologies have a number of limitations. Technological advances, such as genetic engineering and cell culture techniques, can provide cost reduction, increase productivity, and improve ability to respond to emerging biological threats. Next generation platforms such as mRNA and DNA vaccines are paving the way for vaccine development due to their low cost, safety, high efficacy and rapid mass adoption. In addition, the vaccine platform chosen to deliver the antigen(s) of interest may influence the immune response [16].

Although adenoviral vectors are a popular platform for creating vaccines against various flavivirus infections, including those caused by Zika, Dengue, and West Nile viruses, research on TBE is very limited. To date, only two studies have been reported that investigate adenoviruses as candidate vaccines against TBEV [61–63]. In these studies, a vector based on human adenovirus type 5 expressing the non-structural protein NS1 of TBEV was examined. Experiments on mice demonstrated good immunogenicity and protective efficacy; however, no further studies were conducted. Therefore, the recombinant simian adenovirus type 25 expressing prM/E that we developed and characterized is a promising candidate for the development of a TBEV vaccine. Additionally, a similar vector based on the chimpanzee adenovirus ChAdOx1 expressing prM/E of the Zika virus is currently undergoing phase I clinical trials. (Clinical-Trials.gov: NCT04015648 and NCT04440774).

Despite the data described in the scientific literature regarding the phenomenon of antibody-dependent enhancement of infection in vaccinated individuals and those who have recovered from Flavivirus infections, other studies have shown that anti-TBEV immunity induces negligible enhancement of ZIKV infection in a murine model and is unlikely to cause enhanced ZIKV infection in humans [64]. Another study indicates that previous TBEV vaccination does not affect the early IgM-driven neutralizing response to YF17D [65]. Therefore, SAd25 can be considered an effective viral vaccine vector for clinical trials in humans.

Thus, the aim of our work was to evaluate of the immunogenicity and protectivity of candidates TBEV vaccine: chimeric replication-competent attenuated virus YFV 17DD-UN/TBEV(FE), recombinant replication-defective adenovirus SAd25-TBEV(FE) and mRNA-TBEV(FE)-LNP. To generate three candidate vaccines, we used the same antigen – PrM/E of TBEV belonging to Sofjin strain. This made it possible to evaluate the immunogenicity and protective activity of the same antigen depending on the delivery system.

In our previous work [30], we characterized a chimera of YFV 17DD-UN strain and TBEV European subtype, which was made by infectious subgenomic amplicon (ISA) approach. Here we generated chimeric replication-competent attenuated virus YFV 17DD-UN/TBEV(FE) by using bacterial artificial chromosomes. During preliminary experiments we selected dose of virus (not shown). As we expected, the relatively safe dose of the YFV 17DD-UN/TBEV(FE) was lower than for the chimera with the European subtype and amounted to 10^4^ PFU per mouse. This difference is explained by the variability of the PrM/E genes structure of the initial viruses, which determines neurotropism, neurovirulence and neuroinvasiveness of various genetic variants of the TBEV.

According to results of a statistical comparison, mRNA-TBEV(FE)-LNP occupies a leading position in immunogenicity, demonstrating the highest rates of virus-neutralizing antibodies. It is important to notice that 1/10 of the human dose mRNA-TBEV(FE)-LNP (5 μg per mouse) was sufficient to achieve the highest rates of virus-neutralizing antibodies. Whereas, high doses of SAd25 TBEV(Fe) (5×10^10^ virus particles) were used, which is comparable to the human dose of rAD5 in the vaccine Sputnik V (10^11^ viral particles per dose). We did not observe formation of neutralizing antibodies in group mice with 10^4^ PFU of YFV 17DD-UN/TBEV(FE). Experiments with challenge showed that immunization with SAd25 TBEV(Fe) (groups with one and boost immunization) and mRNA-TBEV(FE)-LNP provided reliable protection (100%) with intraperitoneal infection of 100 LD_50_ TBEV. The survival rate of animals was 85,7% in YFV 17DD-UN/TBEV(FE) group. Thus, delivery of PrM/E genes of TBEV in the form of mRNA-LNP is the more promising for the development of a new vaccine. In addition, mRNA-LNP platform allows us to create multi-component cocktails with a wide range of heterologous antigens.

When developing vaccines, it is essential to pay attention not only to the advantages but also to the disadvantages of different antigen delivery system. The ability to maintain the pathogens’ replication potential without causing disease or reversion to virulence is a key consideration for the live attenuated vaccines. The major disadvantage of live attenuated vaccines it’s theirs disease-causing potential in normal and immunocompromised individuals. Addition, this technology requires strict quality control, as well as qualified trained personnel, which leads to increased production costs. Viral vectors are a versatile production platform and in addition to being highly immunogenic, viral vector-based vaccines are easier to produce and, in some cases, safer than inactivated, live attenuated and recombinant protein technologies. The main disadvantages of this platform are pre-existing immunity to the viral vector and reduced effectiveness of subsequent administrations due to antiviral immunity. Compared with viral vaccine platforms, mRNA poses virtually no risk of genomic integration. mRNA vaccines are also more cost-effective and relatively easy to manufacture. Instability at room temperature, dependence on ultra-low cold chain transport, high reactogenicity, and a relatively narrow safety window are the main limitations of the platform. The development of effective and biodegradable lipids and new formulations will likely address the shortcomings of the new platform [17, 66–68].

Our study demonstrates that mRNA based PrM/E antigen delivery elicits a stronger immune response compared to recombinant replication-defective virus or even chimeric replication-competent attenuated virus. Our study has some limitations such as antigenic composition - only one historically known antigen generated from TBEV belonging to Sofjin strain used in our experiments. These limitations are mitigated by the fact that our goal was to compare vaccine platform rather than generate ready to use vaccine candidate Further experiments will allow us to select an antigen composition capable of neutralizing viral isolates of all major genotypes of the virus. Also, the task of further experiments will be to select the immunization regimen and doses, as well as conduct studies on the duration of protection formed by vaccination.

## 5. Conclusions

In this study, we compared candidate TBEV vaccine-candidates based on mRNA, adenovirus serotype 25 and chimera of YFV platforms. Our study demonstrates that the mRNA-based platform is more promising for the creation of a vaccine against TBEV.

## Author Contributions

Conceptualization, N.A.K. and V.A.G.; methodology, N.A.K., E.V.M., A.E.S., D.A.K., M.A.N, O.V.Z., O.P., I.V.V., T.A.O., E.P.M., E.N.B. E.A.M., A.N.Z., E.V.U, A.G.S.; formal analysis, N.A.K., E.V.M., A.E.S., V.I.Z.; investigation, N.A.K., V.A.G., E.V.M., A.E.S., D.A.K., O.V.Z., O.P., I.V.V., T.A.O., E.P.M., E.N.B., E.Y.B., A.N.Z., E.V.S., E.V.U., O.V.U., A.G.S., V.I.Z., D.Y.L., A.L.G.; data curation, N.A.K..; writing—original draft preparation, N.A.K. and V.A.G.; writing—review and editing, N.A.K., V.A.G., A.E.S.; visualization, N.A.K., A.E.S.; supervision, V.A.G.; project administration, V.A.G. All authors have read and agreed to the published version of the manuscript.

## Institutional Review Board Statement

The animal study protocol was approved by the Institutional Animal Care and Use Committee (IACUC) of the Federal Research Centre of Epidemiology and Microbiology named after Honorary Academician N.F. Gamaleya and were performed under Protocol # 63, 05 October 2023.

## Informed Consent Statement

Not applicable.

## Data Availability Statement

The data presented in this study are available on request from the corresponding author.

## Acknowledgments

Many thanks to the employees of Gamaleya Center who provided the purchase of reagents, as well as project support Timofey A. Remizov and Anastasia A. Zakharova.

## Disclaimer/Publisher’s Note

The statements, opinions and data contained in all publications are solely those of the individual author(s) and contributor(s) and not of MDPI and/or the editor(s). MDPI and/or the editor(s) disclaim responsibility for any injury to people or property resulting from any ideas, methods, instructions or products referred to in the content.

